# The Synthesis Success Calculator: Predicting the Rapid Synthesis of DNA Fragments with Machine Learning

**DOI:** 10.1101/2020.06.05.136820

**Authors:** Sean M. Halper, Ayaan Hossain, Howard M. Salis

## Abstract

The synthesis and assembly of long DNA fragments has greatly accelerated synthetic biology and biotechnology research. However, long turnaround times or synthesis failures create unpredictable bottlenecks in the design-build-test-learn cycle. We developed a machine learning model, called the Synthesis Success Calculator, to predict whether a long DNA fragment can be readily synthesized with a short turnaround time. The model also identifies the sequence determinants associated with the synthesis outcome. We trained a random forest classifier using biophysical features and a compiled dataset of 1076 DNA fragment sequences to achieve high predictive performance (F_1_ score of 0.928 on 251 unseen sequences). Feature importance analysis revealed that repetitive DNA sequences were the most important contributor to synthesis failures. We then applied the Synthesis Success Calculator across large sequence datasets and found that 84.9% of the *Escherichia coli* MG1655 genome, but only 34.4% of sampled plasmids in NCBI, could be readily synthesized. Overall, the Synthesis Success Calculator can be applied on its own to prevent synthesis failures or embedded within optimization algorithms to design large genetic systems that can be rapidly synthesized and assembled.

---

DNA synthesis and assembly have become a central technology in modern synthetic biology, enabling the construction of synthetic genetic systems with defined DNA sequences, including recoded genes^1^, engineered proteins^2^, genetic circuits, metabolic pathways and synthetic genomes^3–6^. Whenever a new genetic system does not exist in nature, the synthetic DNA fragments needed to build the genetic system are commonly purchased from commercial service providers. While prices depend on the synthetic DNA fragments’ length and purity as well as other business factors, here we focus solely on the amount of time needed to physically synthesize and assemble them, called the turnaround time. The turnaround time for synthetic DNA fragments is significant as it is often the rate-limiting step in a typical design-build-test-learn cycle for engineering genetic systems^6, 7^. For example, multiple synthetic DNA fragments are commonly needed to build a desired genetic system, and therefore a delay or cancellation in making one DNA fragment will affect the build time of the entire genetic system. Faster and more reliable turnaround times would therefore greatly accelerate the engineering of genetic systems.

When DNA synthesis and assembly takes place normally, turnaround times are currently about 5 to 9 business days for non-clonal DNA fragments (up to 3000 bp) and 11 to 25 business days for purified DNA fragments (up to 5000 bp). However, the presence of several sequence determinants can cause DNA synthesis and assembly to fail, yielding a mixture of low-quality synthetic DNA with several undesired byproducts. In some cases, the process conditions can be adjusted to improve DNA product yields, though with a greatly increased turnaround time. In other cases, the process yields are so low that the requested DNA fragments are never delivered. Therefore, it would be highly beneficial to identify the sequence determinants that interfere with DNA synthesis and assembly, and develop algorithmic approaches to locating them within a desired genetic system’s sequence. Designing genetic systems for rapid synthesis as well as maximal function greatly reduces the turnaround times for building genetic systems.

Here, we developed and validated a machine learning classifier to quantify the effects of several sequence determinants on synthesis outcomes and to accurately predict when a DNA fragment could be synthesized with a short turnaround time. During evaluation, the classifier also identifies the locations of the sequence elements associated with DNA synthesis failure with a quantified priority for removal (**Figure 1A**). To do this, we identified 38 DNA sequence properties that could potentially interfere with DNA synthesis and assembly and compiled a database containing 1076 DNA fragment sequences with known synthesis outcomes. We then developed an automated train-validate-test pipeline to develop a random forest classifier that was both generalized and accurate. Altogether, the finalized random forest classifier, called the Synthesis Success Calculator, achieved an F_1_ score of 0.928 on 251 unseen DNA fragment sequences (**Figure 1B**). Surprisingly, we found only 9 sequence properties associated with a DNA fragment’s synthesis success; depending on their values, there is a high chance of synthesis failure. We demonstrate the potential applications of the Synthesis Success Calculator by recoding recalcitrant protein coding sequences using minimum amounts of synonymous codon optimization, by evaluating how much of the *E. coli* genome can be readily synthesized, and by estimating the percentage of readily synthesized plasmids in the NCBI database.

**Figure 1:**
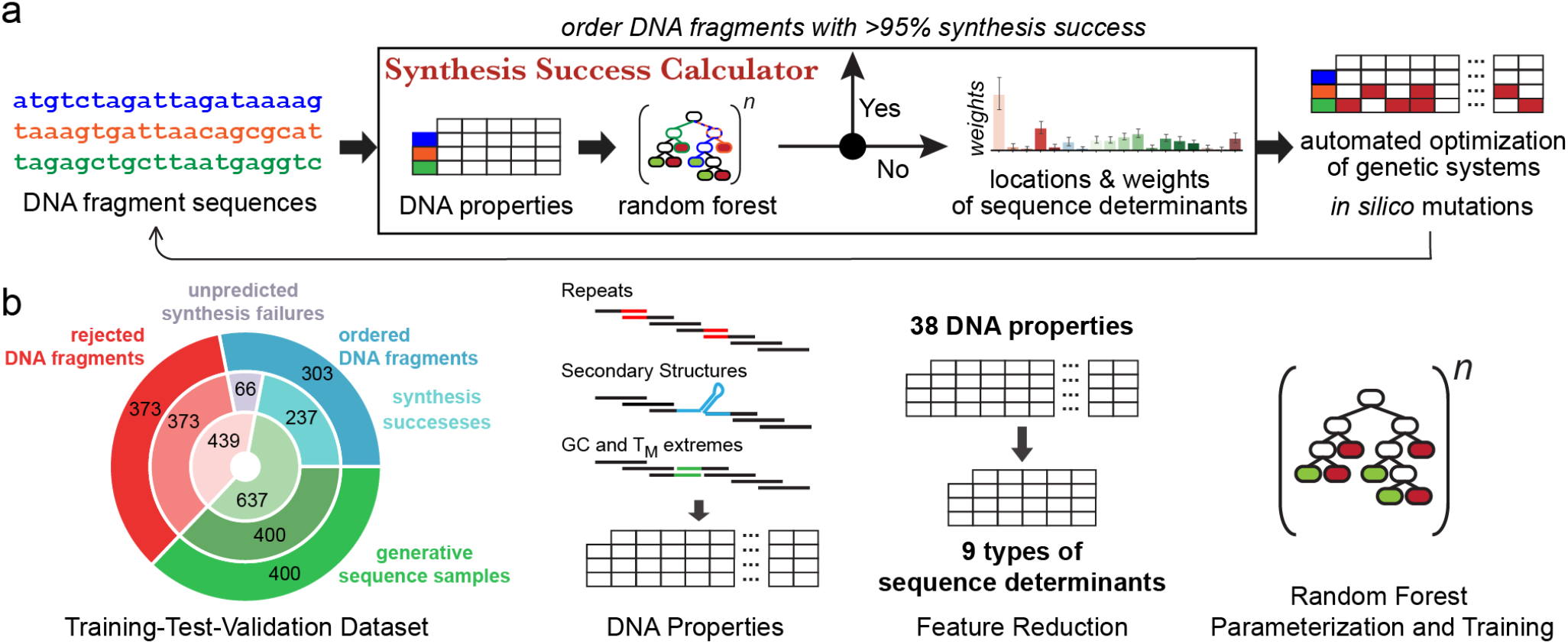
The Synthesis Success Calculator. (a) The Synthesis Success Calculator uses calculated DNA properties and a random forest classifier to predict when a DNA fragment sequence can be synthesized with a fast turnaround time. It also determines the locations and importances of any sequence determinants associated with synthesis failure. Design algorithms can use this information to optimize the genetic system’s sequence for rapid synthesis and maximal function. (b) We created a dataset of 1076 sequences and applied a train-validation-test protocol to parameterize the Synthesis Success Calculator, employing feature reduction to identify the sequence determinants associated with synthesis failure.

Across the synthetic biology ecosystem, algorithms have been developed to break down desired DNA fragments into the optimal set of oligonucleotides for synthesis, for example, Primerize^8^, DNAWorks^9^, Gene2Oligo^10^, and GeneDesign^11^. Other algorithms consider how repetitive sequence elements affect the synthesis and evolutionary robustness of a genetic system^12^, for example, the EFM Calculator^13^. Commercial service providers have also developed proprietary rules for evaluating synthesis complexity, which have been encoded in the Genome Calligrapher^14^ and BOOST^15^ algorithms. However, the Synthesis Success Calculator is distinct from previous efforts as it satisfies several community-wide requirements. First, the Synthesis Success Calculator is fully *transparent* with a non-proprietary dataset, an open-source Python implementation, and a model trained to prioritize researchers’ needs first, for example, by limiting false positives. Second, it includes a pipeline for automatic retraining on any inputted sequence dataset, enabling researchers to identify new sequence determinants affecting synthesis outcomes, for example, when using future chemistries. Third, as a Python module, the Synthesis Success Calculator can be directly embedded within other genetic system design algorithms. To support that capability, the Synthesis Success Calculator also quantifies the sequence determinants’ effects on synthesis success so that design algorithms can more easily find the shared sequence design space where a genetic system achieves both rapid synthesis and maximal function.

## Results and Discussion

We began by enumerating 38 sequence properties that could influence the DNA synthesis and assembly process through a mechanistic interaction (**Supplementary Table 1**). There are 15 properties related to the presence of repetitive sequences with different positions, lengths and densities. Repetitive sequences confound the DNA assembly process by allowing undesired hybridization or annealing interactions, resulting in byproduct formation. There are 7 sequence properties related to the formation of single-stranded secondary structures, which inhibit hybridization between complementary strands during DNA assembly. There are 11 properties related to extreme double-stranded DNA melting temperatures or GC contents, particularly at the terminal ends of DNA fragments, which make it difficult to achieve uniform annealing of oligonucleotides and amplification of full-length DNA fragments. There are 4 properties related to the presence of single-stranded DNA tertiary structures, including G quadruplexes, C-rich i-motifs, and polynucleotide runs. The last property is the DNA fragment’s length.

We then constructed a database of DNA sequences to train, validate, and test the random forest classifier. We first collected 303 DNA fragment sequences with known synthesis success outcomes, taken from our internal order history as well as from contemporary publications that included their synthesized DNA constructs as supplementary information^6, 7^. These DNA fragments were ordered from several different commercial service providers. 66 of these DNA fragments were labeled as synthesis failures because they were either cancelled or significantly delayed beyond the provider’s turnaround time. Notably, even though these DNA fragments passed the commercial service providers’ pre-existing filters for synthesis complexity, there was still a 22% frequency of synthesis failure. The synthesis failure rates are further compounded when multiple DNA fragments were needed to build a genetic system; when ordering 4 DNA fragments, there is only a 37% chance of obtaining them all with a short turnaround time. However, because these DNA fragments did pass the commercial service providers’ filters, they are missing important sequence determinants that are known to create synthesis failure. Therefore, to develop a generalized machine learning model, it becomes necessary to augment the database with a diversified and balanced set of DNA fragment sequences that serve as either positive or negative controls.

As negative controls, we added 373 sequences that were systematically designed to violate the commercial service providers’ filters for high sequence complexity, including the presence of long repetitive DNA sequences, fast folding hairpin structures, palindromic subsequences, polynucleotide runs, sequences with extreme GC content, and sequences with extreme DNA melting temperatures (**Supplementary Table 2**). Sequences in this dataset are rejected by contemporary commercial service providers and therefore are labeled as synthesis failures. As positive controls, we selected 400 subsequences from the successfully synthesized DNA fragments (354 to 1863 base pairs), that each contained a smaller number of individual DNA properties. The purpose of these positive controls is to isolate each DNA property that is associated with synthesis success in order to quantify its importance. Without the negative controls, the random forest classifier could not learn about the DNA properties with known negative effects. Without the positive controls, the random forest classifier would not be able to learn the importance of individual DNA properties. The presence of these controls also focuses the random forest classifier on learning why 22% of the synthesized DNA fragments were not successful. Altogether, the database contains 1076 sequences with a well-balanced combination of evaluated DNA properties and synthesis outcomes (all sequences are listed in **Supplementary Data 1**).

We selected a random forest classifier as our machine learning model for several reasons. Random forest models use an ensemble of decision trees to predict how combinations of properties (“features”) and their values determine an outcome. Compared to other machine learning models, random forests require fewer datapoints for optimal training; here, fewer means hundreds to thousands and not millions of data points. When their hyperparameters are optimized, for example, the number of features used per tree, the number of trees per forest, and the thresholds for creating new branches, random forest models do not over-fit on a dataset (“no memorization”), but their predictions can readily generalize across the dataset, including the unseen test set. Using feature drop analysis, random forest models can also be trained to reject properties that have limited effect on the outcome. Finally, when a random forest predicts an outcome, we can readily visualize and understand how it made that decision, including the locations of the DNA properties in the fragment sequence that are associated with synthesis failure.

We started by developing an automated test-train-validate pipeline commonly used for training machine learning models, utilizing the scikit-learn Python package^16^. We ran three rounds of hyperparameter optimization to develop a baseline random forest classifier, utilizing either randomized or grid search and employing 10-fold cross-validation to measure model performance (**Methods**). Model accuracy is quantified according to its F_1_ score, Matthews Correlation coefficient (MCC) and Cohen’s Kappa score (CK), which together provide a metric of the model’s precision, recall, and information content (**Methods**). Using these baseline hyperparameters, we then carried out a feature reduction strategy, called drop-column, that quantifies the contribution of each DNA property to model accuracy. To do this, the train-validate-test pipeline is carried out anew in the presence versus absence of each individual DNA property, using the same 80-20 train-test split and 100 random initial conditions, followed by comparing the model accuracies (**Figure 2A**). A DNA property’s drop-column importance is the average difference in model accuracy when excluding a DNA property across 100 iterations of the train-test process. Overall, we found that only 9 DNA properties were important enough to lead to a reduction in model performance (**Figure 2B**). Removing any of other 29 DNA properties during the train-test process resulted in a lower F_1_-MCC-CK combined score, for example, by reducing the information content in the model. Here, the most important DNA property was the length of the longest repetitive sequence within the DNA fragment, though overall GC content, extreme double-stranded DNA melting temperatures, GC-rich single-stranded DNA hairpins, and DNA fragment size were also important. These DNA properties are verified as sequence determinants of synthesis success.

**Figure 2.**
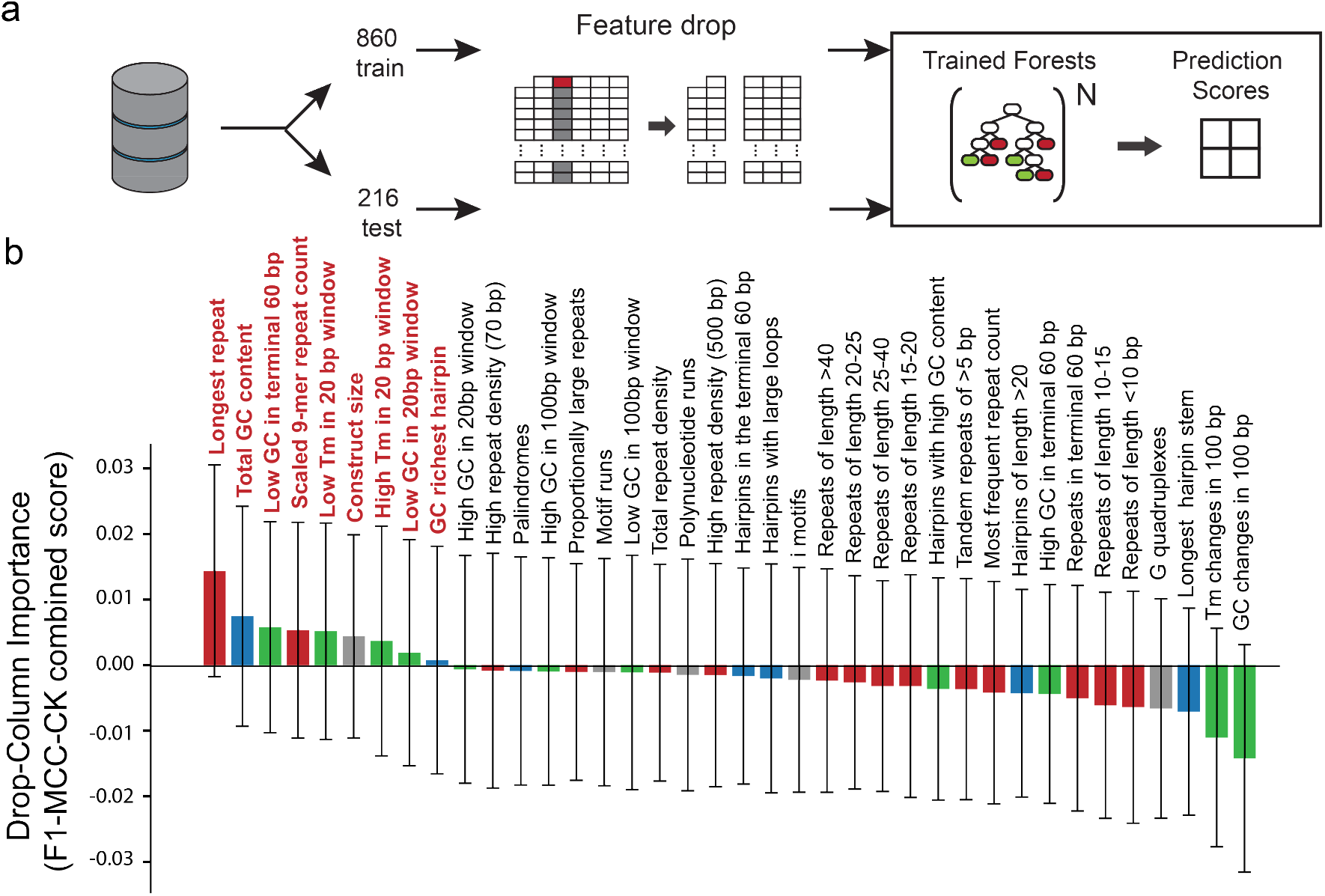
Feature reduction using the 1076 sequence dataset. (a) The workflow used to quantify the importance of each feature in the random forest classifier. (b) Drop column importance of each feature to the classification of 216 test set sequences, colored according to their general categories as repeats (red), secondary structures (blue), GC content (green), or miscellaneous (grey). Error bars represent standard deviations across 100 randomly initialized random forests (n = 100).

We then began a new train-validate-test pipeline to train the random forest model, but now considering only our reduced set of 9 sequence determinants across the 1076 DNA fragment sequences in the database. We carried out training on 40,000 randomly initialized random forest configurations utilizing a training set of 575 DNA fragment sequences, followed by a performance assessment on a validation set of 250 DNA fragment sequences (**Figure 3A**). The resulting random forest classifier had a F_1_ score of 0.928, an MCC of 0.851, and a CK of 0.851 when evaluated on this validation set (**Figure 3B**). We then performed a more rigorous assessment of the classifier’s performance by evaluating its predictions on an unseen test set of 251 DNA fragment sequences, yielding a F_1_ score of 0.908, an MCC of 0.810, and a CK of 0.810 (**Supplementary Figure 1**). However, we noted in this test set that there 13 positive predictions (synthesis success) were labeled as negative outcomes (synthesis failure), yielding a false discovery rate of 8.6%. The false discovery rate is the number of incorrectly predicted synthesis successes (false positives) over the total number of predicted synthesis successes. As our objective is to minimize the volatility of the DNA synthesis and assembly process, we then pursued a rebalancing strategy, called oversampling, to develop a random forest classifier with a lower false discovery rate.

**Figure 3:**
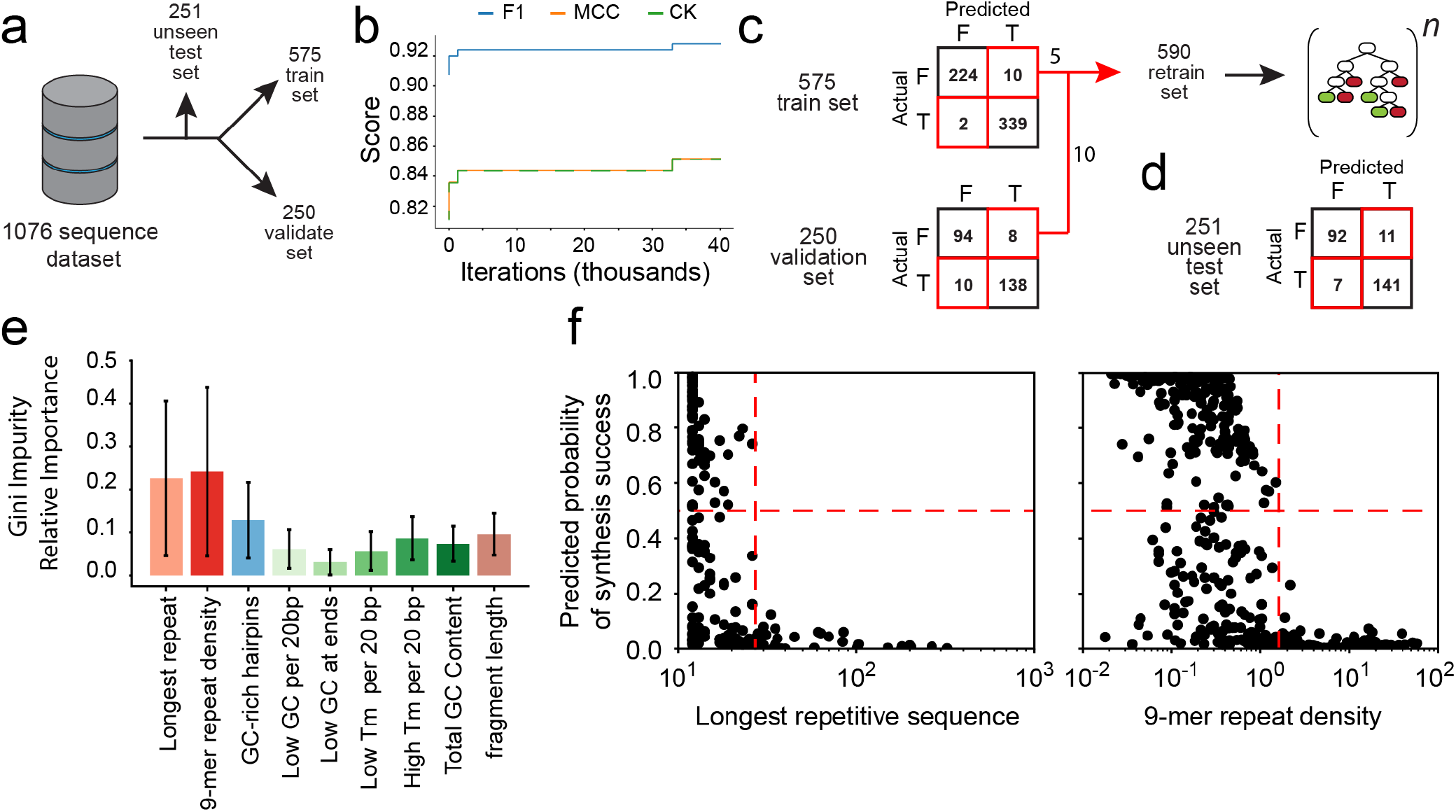
Training and assessment of the Synthesis Success Calculator. (a) A train-validate-test pipeline was used to parameterize a random forest classifier. (b) After hyperparameter optimization, training was performed on 40,000 random forest configurations. F1 is the F_1_ score, MCC is the Matthew’s correlation coefficient, and CK is Cohen’s Kappa. (c) The predicted and actual synthesis outcomes are shown when evaluated on the train and validation sequence sets, using the best random forest configuration from (b). Oversampling is carried out to create a more balanced retrain set. (d) The predicted and actual synthesis outcomes are shown when evaluated on the unseen test set, using the retrained random forest classifier. Synthesis outcomes are either success (T) or delayed/cancelled (F). (e) The relative importance values of each sequence determinant in the retrained random forest classifier are shown. Error bars show the standard deviation of importance values across all trees in the forest (n = 1512). (f) The predicted probability of synthesis success is shown across sequence determinants quantifying the presence of repetitive DNA sequences. Dashed red lines show the decision threshold for synthesis success (horizontal) and the sequence determinant values associated with synthesis failure (vertical).

To apply oversampling, we first identified 5 DNA fragment sequences in the training set that contained sequence determinants that were infrequently present, compared to other values (**Figure 3C**). We also identified 10 DNA fragment sequences in the validation set that contained extreme sequence determinant values that were not well-represented in the training set. We then added these 15 fragment sequences to the training set to create a rebalanced training set with 585 fragment sequences. We then ran additional iterations of training using this rebalanced training set (**Figure 3C**). The resulting random forest classifier had an excellent F_1_ score of 0.928, a MCC of 0.851, and a CK of 0.851 when evaluated on the unseen test set, matching the performance on the validation set and yielding an improved false discovery rate of 7.2% (**Figure 3D**). This finalized random forest classifier was designated version 1.0 of the Synthesis Success Calculator.

Random forest classifiers are not black boxes; after training them, we can analyze the decision trees and sequence determinant cutoffs that lead to a successful classifier (**Methods**). Here, we found that the Synthesis Success Calculator considered the presence of repetitive sequences as the property most associated with synthesis failure (**Figure 3E**). Two sequence determinants – the length of the longest repetitive sequence and the number of repetitive 9-mers per 100 base pairs – had the highest Gini importances, which quantify how frequently they were used to make a decision within the random forests’ decision trees.

We also examined how the Synthesis Success Calculator’s sequence determinant cutoffs controlled its predictions. For example, the Synthesis Success Calculator applied a fairly sharp cut-off to the longest repetitive sequence in a DNA fragment; DNA fragments with a 26 base pair (bp) repetitive sequence were predicted to be readily synthesized, but any repetitive sequence longer than 26 bp would tilt the classifier’s prediction towards synthesis failure (**Figure 3F**). Similarly, if the density of repetitive sequences was too high, the Synthesis Success Calculator would predict synthesis failure; for example, if any 100 bp of the DNA fragment contained 16 or more overlapping repetitive sequences, each 9 bp long (9-mer repeats), then the classifier predicted synthesis failure. Overall, the Synthesis Success Calculator would also predict synthesis success if a DNA fragment’s GC content was between 29% and 63%.

Next, we applied the Synthesis Success Calculator across three illustrative examples and identified how sequence determinants affected the synthesis outcomes, assisted by the Python package treeinterpreter^17, 18^. First, we uniformly broke up the *Escherichia coli* MG1655 genome (NCBI Accession #NC000913.3) into 2320 DNA fragments, each 2500 bp long, and evaluated their synthesis success. According to the classifier, 84.9% of the DNA fragments were predicted to be readily synthesized with a short turnaround time (**Figure 4A**). The remaining DNA fragments (15.1%) were predicted to be difficult to synthesize, due to the presence of single-stranded DNA secondary structures with high GC content, regions with very high double-stranded DNA melting temperatures, and regions with a long repetitive sequence. In the second example, we extracted 4204 unique protein coding sequences from same genome (42 to 7077 bp long) and predicted how readily we could synthesize a DNA fragment containing each coding sequence. The classifier predicted that 84.5% of these DNA fragments could be readily synthesized with a short turnaround time. The classifier identified several failure-associated sequence determinants in the remaining DNA fragments (15.5%), including regions of high double-stranded DNA melting temperature, regions containing GC-rich DNA hairpins, and less importantly, the lengths of the DNA fragment (**Figure 4B**). Interestingly, long repetitive DNA sequences or high densities of repetitive sequences were not predicted to be significant causes of synthesis failure. Overall, we were generally surprised that the classifier deemed *E. coli* genomic DNA to be so readily synthesized; however, these predictions are consistent with recent synthetic *E. coli* genome engineering efforts, where the supermajority of genome-like DNA fragments were synthesized within the anticipated turnaround time.

**Figure 4:**
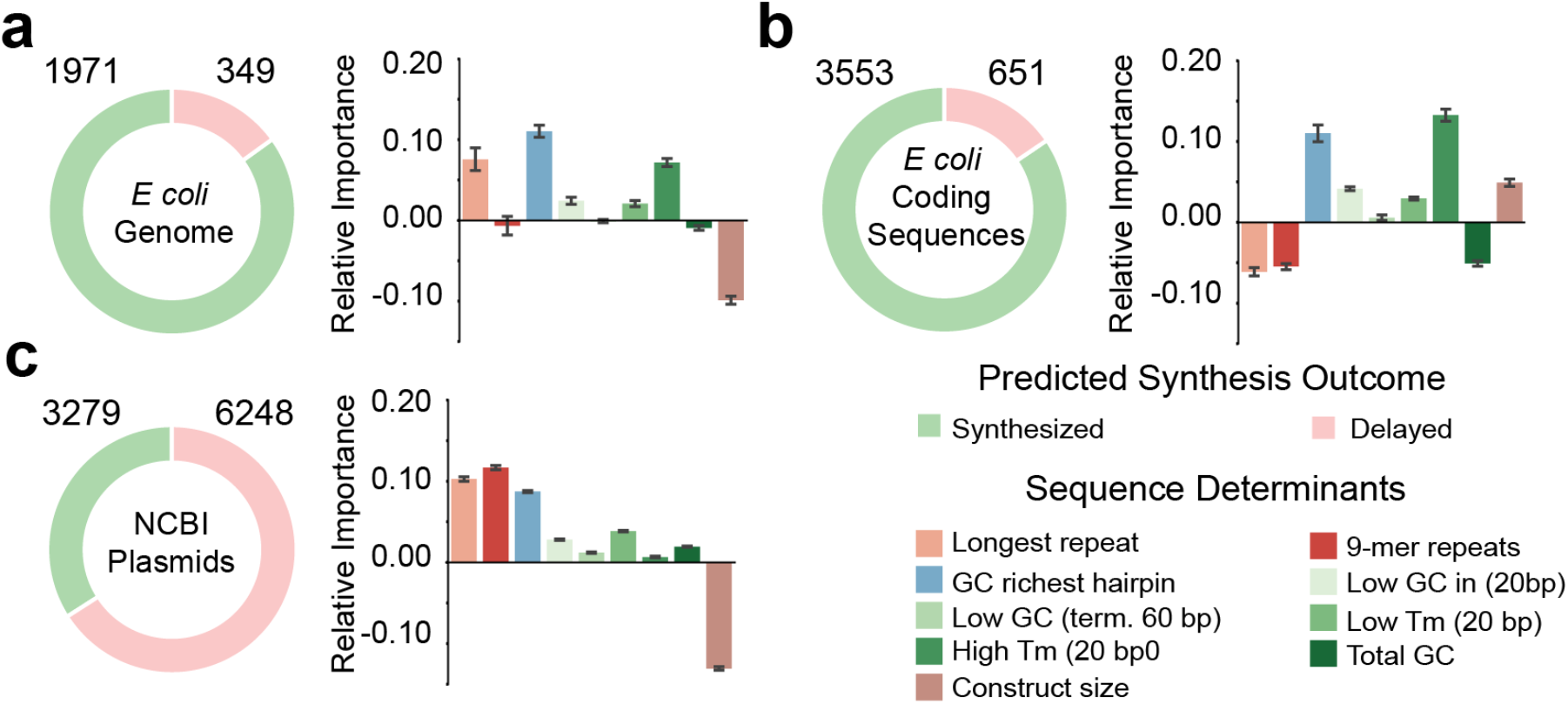
Synthesis Success Calculator Examples. (a) Synthesis success or failure were predicted across 2320 DNA fragments (each 2500 bp) extracted from the *Escherichia coli* MG1655 genome. Considering only the predicted synthesis failures, the relative importance values for each sequence determinant are shown. Error bars represent the standard deviation of sequence determinant values across predicted synthesis failures. (b) The same calculations were performed for 4204 unique protein coding sequences found in the *Escherichia coli* MG1655 genome (42 to 7077 bp long). (c) Synthesis success or failure were predicted across 9527 DNA fragments extracted from 4474 plasmids found in the NCBI database. The color code is shown.

For the third example, we randomly selected 4474 plasmid sequences from the NCBI database, broke them into 9527 DNA fragments (202 to 3500 bp), and applied the Synthesis Success Calculator to predict their synthesis success. In contrast to the previous examples, the classifier predicted that only 34.4% of plasmid sequences could be readily synthesized with a short turnaround time (**Figure 4C**). The classifier predicted that the remaining DNA fragments (65.5%) contained many long repetitive sequences and many regions of high repeat density (9mer repeats) as well as many GC-rich DNA structures. Notably, even though these plasmid-derived DNA fragments have varying lengths, an analysis of the classifier’s predictions showed that fragment length did not negatively impact their synthesis success (**Figure 4C**). All sequences and feature calculations for these three examples are located in **Supplementary Data 2**.

Next, we demonstrated how to embed the Synthesis Success Calculator’s predictions into a design algorithm that optimizes a genetic system’s sequence for improved synthesis success. As an illustrative example, we considered the 651 *E. coli* protein coding sequences that were predicted to be difficult to synthesize (**Figure 4B**) and applied the Synthesis Success Calculator to highlight the most important nucleotide sequence regions associated with synthesis failure. Here, we created a ranked list of these sequence regions, according to the calculated importances of the sequence determinants. We then varied the coding sequences’ synonymous codon usages, but only within these highlighted regions, and at a frequency determined by their importance. Synonymous codon changes were then selected if they improved the Synthesis Success Calculator’s predicted probability of synthesis success (**Methods**). By leveraging the Synthesis Success Calculator’s importance calculations, we found that we only needed to recode 5% of the natural protein coding sequences to push 478 of them (73.4%) into the readily synthesized category. Interestingly, further recoding of the natural coding sequences only provided marginal improvements in predicted synthesis success; recoding 10% or 15% of the natural protein coding sequences resulted in synthesis success rates of 78.3% or 81.4%, respectively (**Supplementary Figure 3**). Overall, the Synthesis Success Calculator was able to pinpoint the most important sequence determinants and identify the sequence regions containing them, enabling a more precise redesign of DNA fragment sequences to ensure their successful synthesis and assembly. All sequences and feature calculations for these designs are located in **Supplementary Data 3**.

Finally, after beginning routine usage of the Synthesis Success Calculator for other Synthetic Biology projects, we observed that our designed DNA fragments had a greater chance of synthesis success. In particular, when multiple genetic parts were encoded within a large DNA fragment, we found that using the Synthesis Success Calculator enabled us to swap out genetic parts that inadvertently created regions with repetitive DNA or extreme melting temperatures. In one example, we designed and ordered 50 synthetic DNA fragments (2859 to 4682 bp long) that each contained multiple promoters, CRISPR RNAs, and reporter proteins, used to build multi-regulator genetic circuits; 42 fragments were successfully synthesized on the first pass (84% success rate), enabling us to immediately proceed with DNA assembly and genome integration. In contrast, collaborators working in parallel on a similar genetic circuit project ordered 18 DNA fragments through the same service provider, but did not use our algorithm for design. Only 3 of the DNA fragments were synthesized and delivered after two attempts (16.6% success rate).

DNA synthesis and assembly are cornerstone technologies in Synthetic Biology, enabling the construction of genetic systems with desired sequences. Regardless of the experimental techniques used, there will always be a set of sequences that remain difficult to synthesize and assemble. Here, we trained and validated the Synthesis Success Calculator, a random forest classifier, that predicts when a DNA fragment can be readily synthesized with a short turnaround time while utilizing contemporary synthesis technologies that have been commercialized, optimized, and scaled-up for higher economies of scale. During that training process, we found that there are only 9 DNA properties associated with synthesis failure with statistical significance, starting from a much longer list of properties and metrics, many of them commonly used by commercial service providers. However, using only those sequence determinants, the Synthesis Success Calculator was able to correctly predict synthesis success with an F_1_ score of 0.928 and a false discovery rate of 7.2% on an unseen test set. Applying the Synthesis Success Calculator, building large genetic systems can now proceed on a more reliable schedule; there is an 74% chance of ordering 4 DNA fragments and receiving them all with a short turnaround time, compared to 37% using pre-existing filters.

To broadly facilitate its usage, we also developed an interactive web interface to the Synthesis Success Calculator, which is available at https://salislab.net/software. After inputting a DNA fragment sequence, the interface reports the predicted outcome (synthesis success or failure), the random forest classifier’s predicted probability of success, and the importance values of the sequence determinants associated with synthesis failure. The interface also highlights the nucleotide regions associated with synthesis failure, illustrating their locations and values using several types of graphs and visual aids.

As new synthesis and assembly techniques are developed, we anticipate that the sequence determinants that control synthesis success will also change, including their cut-off values. We therefore automated the training and testing of the Synthesis Success Calculator, making it easy to input sequence databases, train new random forest classifiers, and extract the important sequence determinants. New DNA properties may also be added to the list of potential sequence determinants. Here, our compiled database contains DNA fragment sequences ordered by several commercial service providers as they utilize similar techniques that all rely on the same DNA properties; nevertheless, it is possible to create separate databases for each distinct synthesis and assembly technology and correspondingly develop separate classifiers. To facilitate this process, the Python source code for the Synthesis Success Calculator is available at https://github.com/hsalis/SalisLabCode.

As demonstrated here, a key purpose of the Synthesis Success Calculator is to enable the automated design of genetic systems for high performance as well as synthesis success. However, when genetic systems are designed using several rules, it can become difficult to find the shared sequence space where all design rules are satisfied, including synthesis success. For example, if a model highlighted too many sequence regions as associated with synthesis failure, then it becomes difficult to modify all those sequences, while retaining the genetic system’s desired functionalities. To overcome this challenge, we restricted the Synthesis Success Calculator to only considering sequence determinants with confirmed associations to synthesis failure, reducing the number of sequence changes needed to ensure synthesis success. To exemplify this distinction, we applied the Synthesis Success Calculator’s predictions on a set of recalcitrant protein coding sequences and found that, for 478 out of 651 proteins, modifying only 5% of their sequences was sufficient to predict synthesis success.

With these confirmed sequence determinants in hand, it is also possible to create well-characterized toolboxes of genetic parts that are specially designed to promote synthesis success. For example, we previously designed and characterized 64 non-repetitive promoters and 28 non-repetitive sgRNA handles, enabling the co-expression of over 20 sgRNAs within Extra Long sgRNA Arrays without introducing repetitive DNA sequences^19^. We also designed and characterized 3500 non-repetitive bacterial promoters, varying transcription rates by over 100,000-fold, that can all be used simultaneously without introducing more than 10 bp repeat sequence^20^. Separately, we designed and characterized another 1722 non-repetitive yeast promoters, varying transcription rates by 25,000-fold, that can all be used simultaneously without introducing more than a 15 bp repeat sequence^20^.

Future design algorithms can combine the Synthesis Success Calculator’s predictions with such synthesis-ready toolboxes of genetic parts to design large genetic systems that have few, if any, sequence determinants associated with synthesis failure. In this way, we can engineer large genetic systems with sophisticated functionalities, while ensuring our ability to synthesize, assemble, and build them.

## Methods

### Software Implementation

The Synthesis Success Calculator was developed in Python 2.7.15 using the Anaconda scientific computing stack^21^. DNA folding predictions were performed using Vienna RNA 2.4.1^22^. Random forests were produced using scikit-learn^16^. Forest analysis and interpretation was done using the treeintepreter^17^ python package and functions from the rfpimp package (https://github.com/parrt/random-forest-importances). External sequences were read and partitioned using Biopython^23^. An implementation of the analysis pipeline used here can be found on GitHub (https://github.com/hsalis/SalisLabCode). The latest version of the Synthesis Success Calculator can be accessed at https://salislab.net/software.

### Dataset Assembly

DNA fragment sequences were sourced from our labs order history over the past 8 years, a recent genome refactoring effort for *S. aureus^6^*, and a recent pressure test to design, build and test metabolic pathways within 90 days^7^. The two external sources were selected due to their explicit, unambiguous discussion of synthesis delays and redesigns.

As the negative controls, DNA fragment sequences were systematically designed to contain DNA properties known to be associated with synthesis failure. To do this, fragment sequences were initialized as non-repetitive, randomized sequences with total GC contents within the acceptable range, followed by purposeful introduction of specific sequence determinants listed in Supplementary Table 2 with varying quantities. These sequences were fed to pre-existing filters hosted by contemporary commercial service providers and selected for inclusion in the negative control set if service providers would not attempt to synthesize it. For the positive controls, we extracted subsequences from readily synthesized DNA fragments with a varied selection of distinct DNA properties, and confirmed that these subsequences also passed pre-existing filters provided by commercial service providers.

### Scoring Metrics

To score the performance of the classifier, we used the geometric mean of the F_1_ score, Matthew’s Correlation Coefficient (MCC), and the Cohen’s Kappa (CK). The F_1_ score measures the harmonic mean of precision and recall, and is defined as

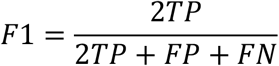

where TP is the true positive rate, FP is the false positive rate, and FN is the false negative rate.

Matthew’s Correlation Coefficient measures the classifier performance, but does so while better accounting for differences in class sizes and is defined as

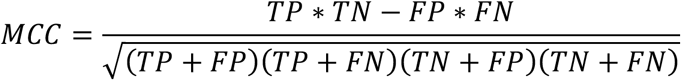

where TN is the true negative rate.

Cohen’s Kappa is a scoring metric that accounts for randomness in the classification by measuring the inter-categorical rating reliability across binary outcomes. It is defined as

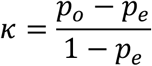

where observed agreement *p*_*o*_ is defined as:

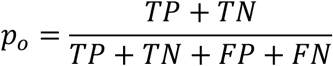

and expected agreement *p*_*e*_ is defined as:

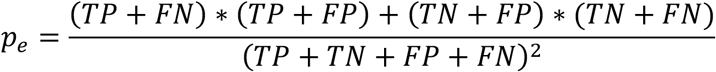

### Model Hyperparameter Optimization

Before training the random forest classifier, we first performed a 3-round iterative hyperparameter optimization, using the full 1076 sequence dataset to find the best hyperparameters to maximize the generalizability of the classifier. We performed iterative 10-fold cross-validation searches for a given set of candidate hyper parameters, using the geometric mean of the F_1_ score, Matthews Correlation coefficient and Cohen’s Kappa as a lumped scoring metric, and took the set of hyperparameters that yielded the best out-of-bag accuracy score from the best performing model of each search.

The first hyperparameter search was 100 iterations of a 200-iteration randomized-search with 10-fold cross-validation, the second was 100 iterations of grid-search with 10-fold cross-validation, and the third search was 200 iterations of grid-search with 10-fold cross-validation, with each subsequent round shrinking the parameter space. During the first round, we varied the number of features seen by each decision tree (max_features), the number of trees in the forest (n_estimators), the number of samples a branch node needs to see in order to split into subsequent nodes (min_samples_split), the number of samples a leaf node needs to receive (min_samples_leaf), and the weight for each class relative to their representation in the dataset (class_weight). Subsequent rounds only varied n_estimators, min_samples_split, and min_samples_leaf (**Supplementary Table 3**).

### Model Feature Reduction

Feature reduction was performed using a 100-iteration drop-column feature importance selection strategy. After one round of hyperparameterization on the full dataset, we split the dataset into an 80-20 train-test split, then scored the model fit for 100 random forests using the set hyperparameters. We then removed one of the features from the dataset, refit the same 100 random forests to the reduced dataset, and calculated their performance scores. We then determined the difference in performance score between using all features versus the reduced 1-feature dropped set. These calculations were repeated for every feature. If the performance score improved when a feature was absent (a negative mean loss), then the feature was removed from the feature set. After feature reduction, 29 DNA properties were removed from the feature set, leaving only 9 DNA properties retained for downstream training of the random forest classifier.

### Model Training and Validation

After hyperparameterization and feature reduction, we trained the random forest using a 575-250-251 train-validate-test split for 40,000 uniquely seeded forests. Class fractions for each sub-dataset were maintained throughout the splits. After the 40,000 iterations, we assessed the performance of the forests on the validation sequences, according to the geometric mean of the F_1_ score, Matthew’s Correlation Coefficient and Cohen’s Kappa, selecting the forest with the highest performance. We then examined the predicted class probability of sequences in the training and validation sets, relative to their known class. Misclassified sequences with extreme outliers in their predicted class properties (greater than 75% for failures, and less than 25% for successes) were added to the training set to create a rebalanced, retraining set. We then carried out additional training of the random forest classifier using the retraining set, containing these oversampled sequences.

### Sequence Determinant Analysis

Sequence determinants that contributed to predicted synthesis failures were extracted using treeinterpreter. The treeinterpreter package calculates the probability prediction of a given tree as

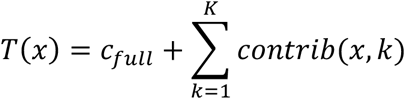

where c_full_ is the default value (bias) of the root node of the tree, K is the number of features, and contrib(x,k) is the contribution of the k^th^ feature in feature vector x along the decision path leading to the leaf node of x. When applied across an entire forest, the probability is stated as

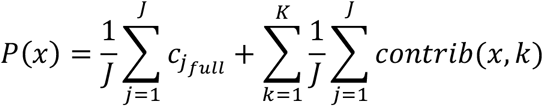

where J is the number of trees in the forest. Using these equations, we quantify the contribution of each sequence determinant to the predicted synthesis outcome.

### Sequence Optimization

To perform targeted recoding on *E. coli* protein coding sequences, we first used treeinterpreter to extract the 3 largest contributing features to the prediction that the sequences were not compatible with rapid synthesis. We then highlighted the nucleotides that contributed to these important features, and allocated different amounts of codon recoding (5%, 10%, 15%, and 20% of the codons) to those highlighted nucleotides. We used a codon table based on the synonymous codon usage of *E. coli* proteins with a high predicted translation initiation rate to recode the highlighted nucleotide regions. After recoding, the coding sequences were reassessed by the classifier to see how well the recoding addressed the predicted synthesis probabilities and whether any of the previously recalcitrant sequences were now predicted to be readily synthesized.

## Author Contributions

SMH and AH compiled sequence databases and implemented algorithms. SMH, AH, and HMS conceived the study and wrote the manuscript.

## Supporting Information

Supplementary Information contains Supplementary Tables 1-3 and Supplementary Figures 1-3. Supplementary Data 1 contains all sequences and feature calculations used to train, test, and validate the Synthesis Success Calculator. Supplementary Data 2 contains DNA sequences and feature calculations to assess the synthesis success of *E. coli* genome segments, *E. coli* coding sequences, and NCBI plasmids. Supplementary Data 3 contains DNA sequences and feature calculations to assess the redesign of protein coding sequences for synthesis success.

## Acknowledgements

This project was supported by funds from the Air Force Office of Scientific Research (FA9550-14-1-0089), the Defense Advanced Research Projects Agency (FA8750-17-C-0254), and the Department of Energy (DE-SC0019090).

## References

1. Patterson, S. S.; Dionisi, H. M.; Gupta, R. K.; Sayler, G. S., Codon optimization of bacterial luciferase (lux) for expression in mammalian cells. J Ind Microbiol Biotechnol 2005, 32 (3), 115–23.

2. Lutz, S., Beyond directed evolution--semi-rational protein engineering and design. Curr Opin Biotechnol 2010, 21 (6), 734–43.

3. Boeke, J. D.; Church, G.; Hessel, A.; Kelley, N. J.; Arkin, A.; Cai, Y.; Carlson, R.; Chakravarti, A.; Cornish, V. W.; Holt, L.; Isaacs, F. J.; Kuiken, T.; Lajoie, M.; Lessor, T.; Lunshof, J.; Maurano, M. T.; Mitchell, L. A.; Rine, J.; Rosser, S.; Sanjana, N. E.; Silver, P. A.; Valle, D.; Wang, H.; Way, J. C.; Yang, L., GENOME ENGINEERING. The Genome Project-Write. Science 2016, 353 (6295), 126–7.

4. Gibson, D. G.; Glass, J. I.; Lartigue, C.; Noskov, V. N.; Chuang, R. Y.; Algire, M. A.; Benders, G. A.; Montague, M. G.; Ma, L.; Moodie, M. M.; Merryman, C.; Vashee, S.; Krishnakumar, R.; Assad-Garcia, N.; Andrews-Pfannkoch, C.; Denisova, E. A.; Young, L.; Qi, Z. Q.; Segall-Shapiro, T. H.; Calvey, C. H.; Parmar, P. P.; Hutchison, C. A., 3rd; Smith, H. O.; Venter, J. C., Creation of a bacterial cell controlled by a chemically synthesized genome. Science 2010, 329(5987), 52–6.

5. Hutchison, C. A., 3rd; Chuang, R. Y.; Noskov, V. N.; Assad-Garcia, N.; Deerinck, T. J.; Ellisman, M. H.; Gill, J.; Kannan, K.; Karas, B. J.; Ma, L.; Pelletier, J. F.; Qi, Z. Q.; Richter, R. A.; Strychalski, E. A.; Sun, L.; Suzuki, Y.; Tsvetanova, B.; Wise, K. S.; Smith, H. O.; Glass, J. I.; Merryman, C.; Gibson, D. G.; Venter, J. C., Design and synthesis of a minimal bacterial genome. Science 2016, 351 (6280), aad6253.

6. Lau, Y. H.; Stirling, F.; Kuo, J.; Karrenbelt, M. A. P.; Chan, Y. A.; Riesselman, A.; Horton, C. A.; Schafer, E.; Lips, D.; Weinstock, M. T.; Gibson, D. G.; Way, J. C.; Silver, P. A., Large-scale recoding of a bacterial genome by iterative recombineering of synthetic DNA. Nucleic Acids Res 2017, 45 (11), 6971–6980.

7. Casini, A.; Chang, F. Y.; Eluere, R.; King, A. M.; Young, E. M.; Dudley, Q. M.; Karim, A.; Pratt, K.; Bristol, C.; Forget, A.; Ghodasara, A.; Warden-Rothman, R.; Gan, R.; Cristofaro, A.; Borujeni, A. E.; Ryu, M. H.; Li, J.; Kwon, Y. C.; Wang, H.; Tatsis, E.; Rodriguez-Lopez, C.; O’Connor, S.; Medema, M. H.; Fischbach, M. A.; Jewett, M. C.; Voigt, C.; Gordon, D. B., A Pressure Test to Make 10 Molecules in 90 Days: External Evaluation of Methods to Engineer Biology. J Am Chem Soc 2018, 140 (12), 4302–4316.

8. Tian, S.; Yesselman, J. D.; Cordero, P.; Das, R., Primerize: automated primer assembly for transcribing non-coding RNA domains. Nucleic Acids Res 2015, 43 (W1), W522–6.

9. Hoover, D. M.; Lubkowski, J., DNAWorks: an automated method for designing oligonucleotides for PCR-based gene synthesis. Nucleic Acids Res 2002, 30 (10), e43.

10. Rouillard, J. M.; Lee, W.; Truan, G.; Gao, X.; Zhou, X.; Gulari, E., Gene2Oligo: oligonucleotide design for in vitro gene synthesis. Nucleic Acids Res 2004, 32 (Web Server issue), W176–80.

11. Richardson, S. M.; Olson, B. S.; Dymond, J. S.; Burns, R.; Chandrasegaran, S.; Boeke, J. D.; Shehu, A.; Bader, J. S., Automated Design of Assemblable, Modular, Synthetic Chromosomes. Parallel Processing and Applied Mathematics, Part Ii 2010, 6068, 280–+.

12. Tang, N. C.; Chilkoti, A., Combinatorial codon scrambling enables scalable gene synthesis and amplification of repetitive proteins. Nat Mater 2016, 15 (4), 419–24.

13. Jack, B. R.; Leonard, S. P.; Mishler, D. M.; Renda, B. A.; Leon, D.; Suarez, G. A.; Barrick, J. E., Predicting the Genetic Stability of Engineered DNA Sequences with the EFM Calculator. ACS Synth Biol 2015, 4 (8), 939–43.

14. Christen, M.; Deutsch, S.; Christen, B., Genome Calligrapher: A Web Tool for Refactoring Bacterial Genome Sequences for de Novo DNA Synthesis. ACS Synth Biol 2015, 4 (8), 927–34.

15. Oberortner, E.; Cheng, J. F.; Hillson, N. J.; Deutsch, S., Streamlining the Design-to-Build Transition with Build-Optimization Software Tools. ACS Synth Biol 2017, 6 (3), 485–496.

16. Pedregosa, F.; Varoquaux, G.; Gramfort, A.; Michel, V.; Thirion, B.; Grisel, O.; Blondel, M.; Prettenhofer, P.; Weiss, R.; Dubourg, V., Scikit-learn: Machine learning in Python. Journal of machine learning research 2011, 12 (Oct), 2825–2830.

17. Saabas, A. Interpreting random forests. http://blog.datadive,net/interpreting-random-forests/ (accessed March 5th).

18. Palczewska, A.; Palczewski, J.; Robinson, R. M.; Neagu, D., Interpreting random forest models using a feature contribution method. 2013, 112–119.

19. Reis, A. C.; Halper, S. M.; Vezeau, G. E.; Cetnar, D. P.; Hossain, A.; Clauer, P. R.; Salis, H. M., Simultaneous repression of multiple bacterial genes using nonrepetitive extra-long sgRNA arrays. Nature biotechnology 2019, 37 (11), 1294–1301.

20. Hossain, A.; Halper, S. M.; Cetnar, D. P.; Reis, A. C.; Salis, H. M., Automated Design of Thousands of Non-Repetitive Parts for Engineering Stable Genetic Systems. Nature biotechnology, accepted.

21. Grüning, B.; Dale, R.; Sjödin, A.; Chapman, B. A.; Rowe, J.; Tomkins-Tinch, C. H.; Valieris, R.; Köster, J.; The Bioconda, T., Bioconda: sustainable and comprehensive software distribution for the life sciences. Nature Methods 2018, 15 (7), 475–476.

22. Lorenz, R.; Bernhart, S. H.; Honer Zu Siederdissen, C.; Tafer, H.; Flamm, C.; Stadler, P. F.; Hofacker, I. L., ViennaRNA Package 2.0. Algorithms Mol Biol 2011, 6, 26.

23. Cock, P. J.; Antao, T.; Chang, J. T.; Chapman, B. A.; Cox, C. J.; Dalke, A.; Friedberg, I.; Hamelryck, T.; Kauff, F.; Wilczynski, B.; de Hoon, M. J., Biopython: freely available Python tools for computational molecular biology and bioinformatics. Bioinformatics 2009, 25 (11), 1422–3.

